# *Candida albicans* filamentation does not require the cAMP-PKA pathway in vivo

**DOI:** 10.1101/2022.03.25.485899

**Authors:** Rohan S. Wakade, Juraj Kamara, Melanie Wellington, Damian J. Krysan

## Abstract

*Candida albicans* is one of the most prevalent human fungal pathogens. Its ability to transition between budding yeast and filamentous morphological forms (pseudohyphae and hyphae) is tightly associated with its pathogenesis. Based on in vitro studies, the cAMP-Protein Kinase A (PKA) pathway is a key regulator of *C. albicans* morphogenesis. Using an intravital imaging approach, we investigated the role of the cAMP-PKA pathway during infection. Consistent with their roles in vitro, the downstream effectors of the cAMP-PKA pathway Efg1 and Nrg1 function, respectively, as an activator and a repressor of in vivo filamentation. Surprisingly, strains lacking the adenylyl cyclase, *CYR1*, showed only slightly reduced filamentation in vivo despite being completely unable to filament in RPMI+10% serum at 37°C. Consistent with these findings, deletion of the catalytic subunits of PKA (Tpk1 and Tpk2), either singly or in combination, generated strains that also filamented in vivo but not in vitro. In vivo transcription profiling of *C. albicans* isolated from both ear and kidney tissue showed that the expression of a set of 184 environmentally responsive correlated well with in vitro filamentation (R^2^ 0.62-0.68) genes. This concordance suggests that the in vivo and in vitro transcriptional responses are similar but that the upstream regulatory mechanisms are distinct. As such, these data emphatically emphasize that *C. albicans* filamentation is a complex phenotype that occurs in different environments through an intricate network of distinct regulatory mechanisms.

**Importance:** The fungus *Candida albicans* causes a wide range of disease in humans from common diaper rash to life-threatening infections in patients with compromised immune systems. As such, the mechanisms for its ability to cause disease are of wide interest. An intensely studied virulence property of *C. albicans* is its ability to switch from a round yeast form to filament-like forms (hyphae and pseudohyphae). Surprisingly, we have found that a key signaling pathway that regulates this transition in vitro, the protein kinase A pathway, is not required for filamentation during infection of the host. Our work not only demonstrates that the regulation of filamentation depends upon the specific environment *C. albicans* inhabits but also underscores the importance of studying these mechanisms during infection.

---

*Candida albicans* is the most common human fungal pathogen and causes disease in both immunocompetent and immunocompromised patients (1). Accordingly, the mechanisms of *C. albicans* pathogenesis have been of broad and ongoing interest (2). One of the most intensively studied *C. albicans* virulence traits is its ability to transition between yeast and filamentous (hyphae and pseudohyphae) morphologies (3). All three morphotypes are observed in infected tissue by histology and strains that are genetically “locked” as yeast or filaments have reduced ability to cause disease, indicating that the ability to transition between morphologies is important for pathogenesis (4).

Consistent with its importance to *C. albicans* pathobiology, the genetic and molecular mechanisms of filamentation have been extensively studied and a large set of genes positively and negatively affect the process (1, 5). Of the signaling pathways involved, the cAMP-Protein Kinase A (PKA) pathway (Fig. 1A) has been shown to regulate in vitro filamentation in response to a wide range of stimuli including heat, carbon dioxide, and bacterial glycopeptides (6). Previous studies also support a model whereby the PKA pathway regulates filamentation through the direct phosphorylation/activation of Efg1 (7) and the indirect de-activation of Nrg1, a transcriptional repressor of filamentation (8). Efg1 is a bHLH type transcription factor that is required for filamentation in most but not all in vitro conditions (9). Efg1 contains multiple candidate PKA phosphorylation sites; mutation of one of these, threonine 206, to a non-phosphorylatable alanine (*efg1*^*T206A*^) reduces filamentation in Spider and GlcNAc medium but not in the presence of serum (7). Overexpression of *EFG1* also partially suppresses the filamentation defect of a strain lacking one of the catalytic subunits of PKA (Tpk2). Nrg1 is a conserved transcriptional repressor that must be de-repressed or degraded for filamentation to occur and strains lacking *NRG1* are constituently filamentous (8). The regulation of *NRG1* expression/degradation has also been genetically linked to the PKA pathway. Although the role of the PKA-Efg1-Nrg1 axis has been well-established in vitro, its effect on filamentation during mammalian infection has not been studied directly.

**Figure 1.**
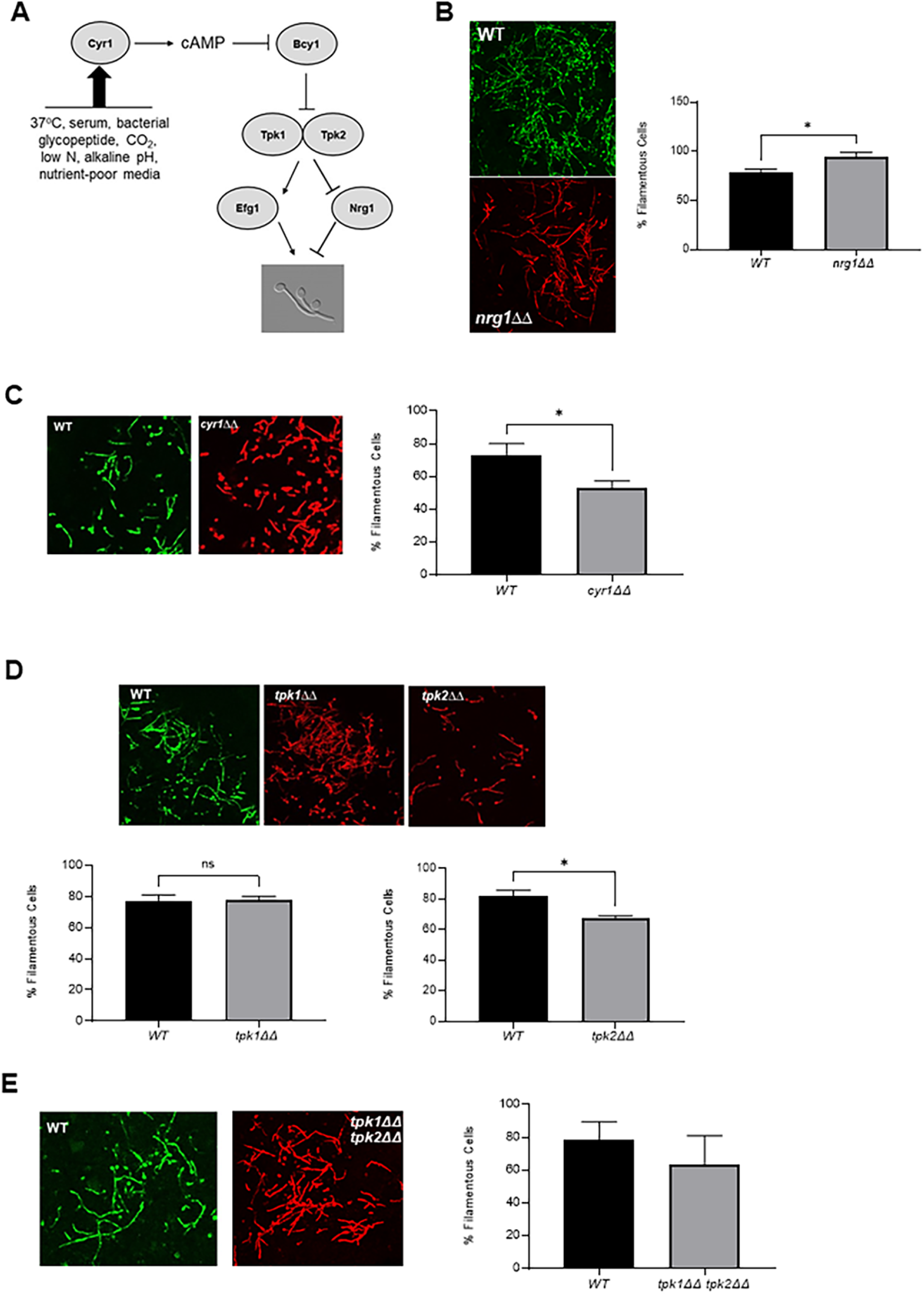
The cAMP-protein kinase A pathway is not required for *C. albicans* filamentation in vivo. A. Simplified schematic of the cAMP-protein kinase A pathway and its in vitro inducers. B-D. show images of WT (green) and the indicated mutant (red) visualized by confocal microscopy visualized 24 hr after co-inoculation into the pinna of a DBA/2 mouse. The accompanying graphs show quantitation of the percent of filamentous cells using the scoring system described in the Materials and methods section. The bars represent at least two independent replicates in which >100 cells were scored in multiple fields. The error bars indicate standard deviation and asterisks indicate that WT and the mutant differ in a statistically significant amount (Student’s t test, p<0.05). B) *nrg1*ΔΔ mutant; C) *cyr1*ΔΔ mutant; D) *tpk1*ΔΔ, *tpk2*ΔΔ, and *tpk1*ΔΔ *tpk2*ΔΔ mutants.

Recently, we developed an intravital imaging approach to the morphological characterization of *C. albicans* in infected tissue by directly injecting fluorescently labeled *C. albicans* into the subepithelial/submucosal tissue of mice ears (10, 11). The *C. albicans* infection cycle begins with the yeast penetrating through the mucosal/epithelial cells to the submucosal/stromal tissue from which it then gains access to the vasculature and disseminates (2). The ear model directly establishes infection within the stroma and thus recapitulates this intermediate step of the cycle (10, 11). In this way, the distribution of yeast and filamentous forms of the fungus during infection of a pathogenically relevant tissue can be quantitatively characterized.

As we reported previously (11), Efg1 is required for filamentation in this model in both reference strains and clinical isolates. Consequently, we were interested to determine if other components of the cAMP-PKA pathway (Fig. 1C) also regulated in vivo filamentation. To do so, a 1:1 mixture of NEON-labeled WT reference and an i-RFP-labeled homozygous deletion mutant was injected into the subdermal ear tissue of a DBA/2 mouse; 24 hr post-infection, the mouse ear was imaged by confocal microscopy and the ratio of filamentous and yeast cells determined for WT and the mutant (see materials and methods for details). Consistent with our previous reported data (11), WT cells show a consistent percentage of filaments (80%) 24 hours post infection; this percentage is similar to that observed in vitro after cells are induced with RPMI+10% serum at 37°C for 4 hr (Fig. 1B). In vitro, deletion of *NRG1* leads to a constitutively hyphal strain (12, 13). Consistent with the in vitro phenotype, the *nrg1*ΔΔ mutant also formed an increased percentage of hyphal cells in vivo (Fig. 1B). Taken together with our previously reported results for the *efg1*ΔΔ mutant, these data indicate that the downstream transcriptional regulators of the PKA pathway carry out similar functions in vitro and in vivo.

In vitro, the adenylyl cyclase, *CYR1*, integrates hypha-inducing inputs from multiple stimuli and pathways to activate the PKA pathway by generating cAMP (Fig. 1A, ref. 3&6). Surprisingly, a strain lacking *CYR1* formed filaments in vivo nearly as well as WT (Fig. 1C) while no hyphae were formed in vitro (Fig. S1A), indicating that cAMP-independent pathways predominately regulate filamentation in vivo. Parrino et al. (14) and Min et al. (15) have described cAMP-independent filamentation in vitro based on the isolation of extragenic suppressors of the *cyr1*ΔΔ mutant that are able to filament in vitro. Characterization of these suppressors revealed that the suppression was due to loss of function mutations in *BCY1*, the gene that encodes the cAMP-responsive negative regulator of PKA (Fig. 1A). If this bypass mechanism were operative in vivo, then filamentation would still be dependent on the PKA kinase isoforms, *TPK1* and *TPK2*. As shown in Fig. 1D, deletion of neither PKA isoform reduces filamentation dramatically in vivo while in RPMI+10% serum both *tpk1*Δ and *tpk2*Δ mutants have modest defects (Fig. S1B).

It remained possible that Tpk1 and Tpk2 have redundant roles during in vivo filamentation. Recently, strains lacking both Tpk1 and Tpk2 subunits have been generated and shown to be unable to filament in vitro (16). To test the Tpk1/2 dependence of in vivo filamentation, we constructed an iRFP-tagged *tpk1*ΔΔ *tpk2*ΔΔ mutant and confirmed that it was unable to filament in vitro (Fig. S1C). In contrast to in vitro conditions, and consistent with the results with the *cyr1*ΔΔ mutant, the *tpk1*ΔΔ *tpk2*ΔΔ double mutant forms filaments in vivo (Fig. 1E). Taken together these data indicate that three of the key components of the cAMP-PKA pathway are largely dispensable for filamentation in vivo.

Since the transcriptional regulators of the canonical cAMP-PKA pathway retain their roles in vitro and in vivo, we predicted that the transcriptional responses during filamentation in vitro and in vivo would also be similar. To test this prediction, we compared the expression profile of our reference strain (SN425) during yeast phase growth (30°C, YPD) to in vitro induction with RPMI+10% BCS at 37°C and to cells isolated from two sites of in vivo filamentation (ear tissue and kidney tissue following disseminated infection). We used a Nanostring probe set (185 genes) based on one previously used by Xu et al. (17) to profile *C. albicans* isolated from kidney organs following disseminated infection. This set contains environmentally responsive genes of which 25% are hyphae-associated (for full list see Table S1).

Cells were isolated after 4 hr of induction while the in vivo samples were collected at 24 hr post-infection; the same time points used form morphological characterization. The absolute expression of the genes was significantly correlated for all three samples. The correlation between expression in the kidney and ear (Fig 2A) was slightly stronger than either infection site with in vitro induction conditions (Fig 2B/C). Although this is a small set of genes, the strong correlation in expression between *C. albicans* in the ear and the kidney supports the notion that filamentation in the ear is transcriptionally similar to the kidney and, although there are likely to be some differences, the two sites are reasonably comparable. The similarity of the transcriptional responses is further supported by the fact that very few genes are uniquely regulated in a single condition (Fig. 2D). Thus, the cAMP-PKA independence of in vivo filamentation cannot be explained by dramatic differences in the transcriptional responses and indicate upstream regulators of the transcriptional responses are distinct between in vitro induction and in vivo infection of tissue.

**Figure 2.**
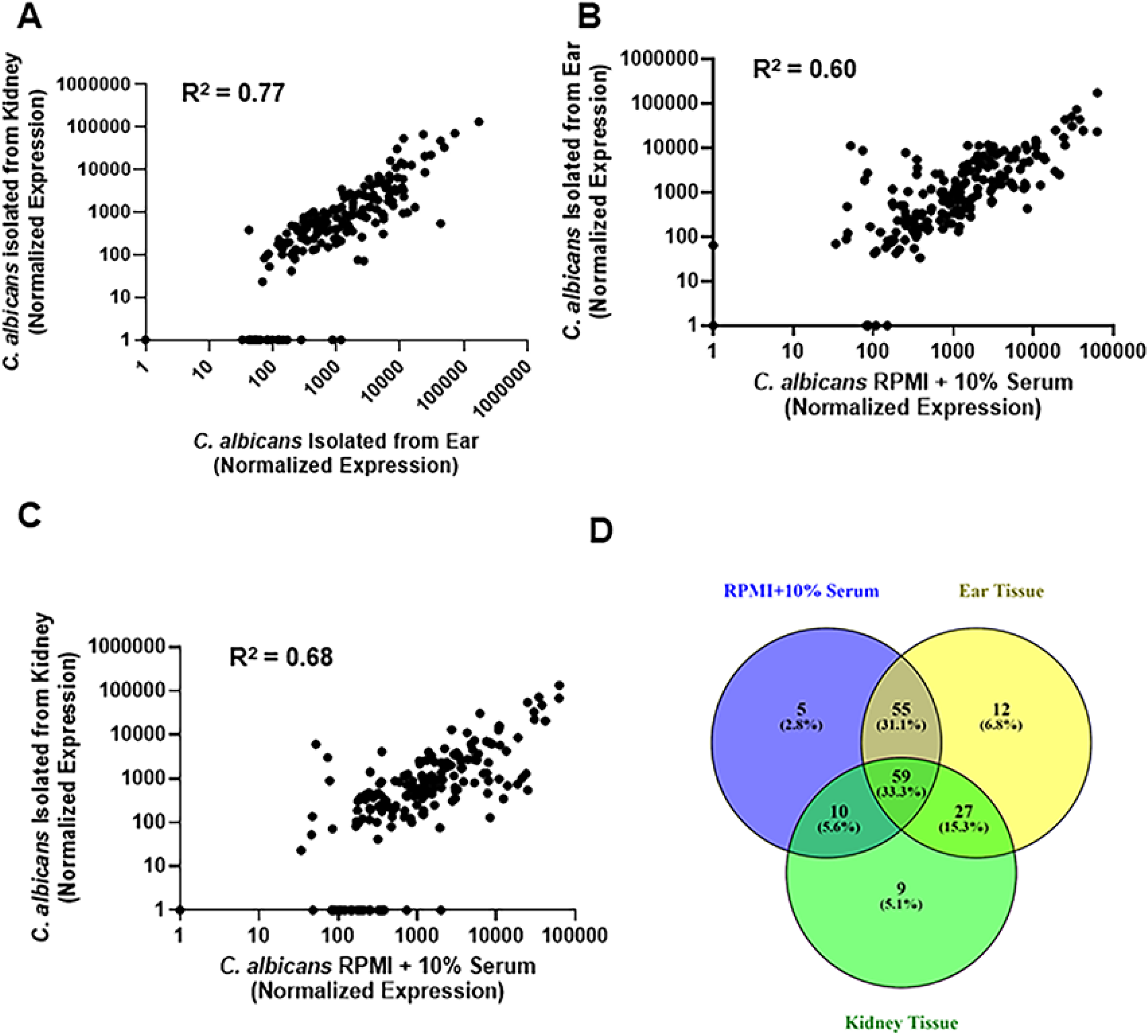
The expression of a set of 185 environmentally responsive genes is similar between in vitro hyphae induction and infection of either mouse ear or kidney. The normalized RNA counts measured by Nanostring nCounter for *C. albicans* cells induced to form hyphae in vitro (RPMI+10% serum, 37°C, 4 hr,); isolated from infected ear; isolated from infected kidney were plotted and analyzed for correlation using Pearson’s coefficient. All correlations are statistically significant. A. kidney vs. ear; B. ear vs. in vitro; C. kidney vs. in vitro. D. Venn diagram for comparing the overlap of differentially expressed genes (2-fold change, statistically significant by Student’s t test, p <0.05) between the three conditions. Data for the plots are summarized in Table S1.

These data have important implications for current models of *C. albicans* morphogenesis. First, many studies have shown that in vitro filamentation is largely cAMP-PKA-dependent (3, 5, 6), our data indicate that, while the cAMP-PKA pathway does contribute to filamentation in vivo, cAMP-PKA independent mechanisms appear to predominate. Second, the standard models of *C. albicans* filamentation posit that PKA directly phosphorylates Efg1 to mediate filamentation (7). In vivo, the discordance between the phenotypes of Efg1 and PKA pathway mutants indicates that Efg1 is independent of the cAMP-PKA pathway under these conditions. Third, the consistent roles of Efg1 and Nrg1 under the two conditions, along with the similarity of the transcriptional responses of in vitro and in vivo filamentation, indicate that distinct upstream regulatory pathways mediate a relatively conserved transcriptional response. It must be noted, however, that histological analyses indicate that Efg1 does not appear to be required for filamentation in the oral cavity and, thus, additional Efg1-independent pathways are also likely to be present (18, 19). Overall, these data and previously reported results emphatically emphasize that *C. albicans* filamentation is a complex phenotype that occurs in different environmental niches through an intricate network of distinct regulatory mechanisms.

## Materials and methods

### Strains, cultivation conditions, and media

All *C. albicans* strains were constructed in the SN background (20) except for the *cyr1*ΔΔ mutant which was generously provided by Dr. Jamie Konopka which was constructed in the BWP17 background (14, 15). The *nrg1*ΔΔ deletion was from the collection of Homann et al. (20), obtained from the Fungal Genetics Stock Center, and confirmed by phenotype and genotype. The *tpk1*ΔΔ and *tpk2*ΔΔ were generated using standard CRISPR-based methods. The *tpk1*ΔΔ *tpk2*ΔΔ double mutants were constructed using the recently described CRIME methodology (21). Briefly, both copies of *TPK2* were knocked out by co-transforming the recipient strain with a split *LEU2* cassette with *TPK2* homology at both ends and two *TPK2* gene targeting guide RNAs (oligos used to construct the deletion mutants are listed in Table S2). In the resulting *tpk2ΔΔ* mutants the *LEU2* cassette was recycled by co-transformation of the mutant strain with a *LEU2* and *ARG4* targeting constructs along with an *ARG4* cassette which served as a selection marker. The resulting *tpk2ΔΔ* mutants with excised *LEU2* cassette were then used to construct double *tpk1ΔΔ tpk2ΔΔ* knockout mutants using the same approach as above. Correct integration and absence of the targeted ORFs was confirmed by PCR analysis.

References strains SN and BWP17 were labelled by transforming with pENO1-NEON-NAT1 while mutant strains were transformed with pENO1-iRFP-NAT1 as previously described. All *Candida albicans* strains were pre-cultured over-night in yeast peptone dextrose (YPD) medium at 30^0^C. Standard recipes were used to prepare media (18). RPMI medium was purchased and supplemented with bovine serum (10% v/v). The yeast phase cells for Nanostring analysis were inoculated into fresh YPD medium and incubated for 4 hrs to achieve mid-log phase. For in vitro hyphae induction, *C. albicans* strains were incubated overnight in YPD at 30°C, harvested, and diluted into RPMI + 10% serum at 1:50 ratio and incubated at 37^0^ C for 4 hours. Cells were fixed with formalin, collected and examined by light microscopy.

### Intravital Imaging and scoring

The inoculation and imaging were carried out exactly as described in (11). Filamentous cells had identifiable mother cells and the filamentous projection was at least twice the length of the mother cell body. Yeast cells were round and/or budded cells. Filamentous cells were confirmed by manually following the hyphal projection through each Z stacks (30 stacks). Yeast cells were further required *not* to project through multiple Z stacks. At least 100 cells in multiple fields were scored; experiments were performed in duplicate or triplicate using independent isolates of strains. Statistical significance was determined by the unpaired Student’s t test. The data sets did not show a detectable difference from normality using the Shapiro-Wilk test (*P*>0.05). Statistical tests were performed using GraphPad Prism software.

### In vitro and in vivo Nanostring analysis

For *in vitro* RNA extraction, three independent cultures were grown overnight in YPD at 30°C, harvested, and diluted into RPMI + 10% serum at 1:50 ratio and incubated at 37^0^ C for 4 hours. Cells were collected, centrifuged for 2 min at 11K rpm at RT and RNA was extracted from the pellet according to the manufacturer protocol (MasterPure Yeast RNA Purification Kit, Cat. No. MPY03100). Extraction of RNA from mouse tissue was performed as reported by Xu et al. with some modification (17). Briefly, to extract RNA from mice ear, following procedure was used. After 24 hr post-injection, mouse was euthanized following the protocol approved by the University of Iowa IACUC. The *C. albicans*-infected mouse ear was removed and directly placed in the ice-cold RNA later solution, then the ear was transferred to the mortal and flash frozen with liquid nitrogen. Further samples were ground to a fine powder, which was transferred to 5 ml centrifuge tube and 1 ml of ice-cold Trizol was added. The samples were placed on a rocker at RT for 15 min and then centrifuged at 10K rpm at 4^0^C for 10 min to remove the debris. Cleared Trizol was transferred to 1.5ml Eppendorf tube and 200 µl of RNase free chloroform was added to each sample. Tubes were shaken vigorously for 15 s and kept at RT for 5 min followed by centrifuge at 12K rpm for 15 min at 4^0^C. The cleared aqueous layer was then transferred to new 1.5ml tube and RNA was further extracted according the Qiagen RNeasy kit protocol.

RNA (40 ng for in vitro and 1.4 µg for in vivo) was added to a NanoString codeset mix (Table S1) and incubated at 65^0^C for 18 hours. After hybridization reaction, samples were proceeded to nCounter prep station and samples were scanned on an nCounter digital analyzer. NCounter .RCC files for each sample were imported into nSolver software to evaluate the quality control metrics. Using the negative control probes the background values were first assessed. The mean plus standard deviation of negative control probes value was defined and used as a background threshold and this value is subtracted from the raw counts. The background subtracted total raw RNA counts were normalized against the highest total counts from the biological triplicates. The statistical significance of changes in counts was determined by two tailed Student’s t test (p <0.05). The expression data are summarized in Table S1.

Probes that were below background were set to a value of 1 to allow statistical analysis. The raw counts, normalized counts, and statistical analyses are also provided in Table S1.

## Acknowledgements

This work was supported by NIH grants R01AI133409 (DJK) and R21AI157341 (DJK). We thank Jamie Konopka (Stony-Brook), Aaron Mitchell (Georgia), and Scott Filler (UCLA) for reading early versions of the manuscript and helpful suggestions. Jamie Konopka for providing strains and Robert Wheeler (Maine) for providing plasmids.

## Author Contributions

Conceptualization: Damian J. Krysan, Melanie Wellington

Formal analysis: Rohan W. Wakade, Juraj Kamara, Melanie Wellington, and Damian J. Krysan

Investigation: Rohan S. Wakade, Juraj Kamara

Methodology: Rohan S. Wakade, Juraj Kamara

Supervision: Melanie Wellington, Damian J. Krysan

Writing-original draft: Damian J. Krysan

Writing-review and editing: Damian J. Krysan

## Figure Legends

**Figure S1. In vitro filamentation phenotypes for strains evaluated in vivo**. A-C show images of WT and the indicated mutants after hyphae induction with RPMI+10% serum at 37°C for 4hr. The graphs show the percentage of filamentous cells and represent 2-3 independent experiments in which >100 cells were scored. The error bars indicate standard deviation and asterisks indicate that WT and the mutant differ in a statistically significant amount (Student’s t test, p<0.05).

**Table S1. Summary of in vitro and in vivo expression of a set of 185 environmentally responsive genes**. Raw RNA counts for each gene; normalized RNA counts; fold-change relative to yeast phase growth; statistical significance of change (Student’s t test, p < 0.05) are shown. Fold changes shown in green are genes with statistically significant increased expression of >2-fold. Fold change in red are statistically significant reductions in gene expression of >2-fold.

**Table S2. Oligos used to construct TPK mutants**.

